# The evolutionary stability of plant antagonistic facilitation across environmental gradients and its ecological consequences: soil resource engineering as a case study

**DOI:** 10.1101/2023.02.05.527181

**Authors:** Ciro Cabal, Gabriel A. Maciel, Ricardo Martinez-Garcia

**Affiliations:** High Meadows Environmental Institute, Princeton University, 08544 Princeton, USA; Department of Biogeography and Global Change, National Museum of Natural Sciences, MNCN, CSIC, 28006 Madrid, Spain; ICTP-South American Institute for Fundamental Research - Instituto de Física Teórica da UNESP, Rua Dr. Bento Teobaldo Ferraz 271, 01140-070 São Paulo SP, Brazil; Center for Advanced Systems Understanding (CASUS), Helmholtz-Zentrum Dresden Rossendorf (HZDR), Görlitz, Germany

**Keywords:** ecosystem engineers, facilitation, primary succession, stress gradient hypothesis, soil amelioration, root competition

## Abstract

1. Plant interactions, understood as the net effect of an individual on the fitness of a neighbor, vary in strength and can shift from negative to positive as the environmental conditions change in time and space. Evolutionary theory questions the stability of non-reciprocal interactions in which one plant has a positive net effect on a neighbor, which in return has a negative net impact on its benefactor. This type of interaction is known as antagonistic facilitation.
2. We develop a spatially explicit consumer-resource model for below-ground plant competition, including plants able to mine resources and make them available for any other plant in the community, termed ecosystem engineers. We use the model to assess whether and under which environmental conditions antagonistic facilitation via soil resource engineering is evolutionarily stable.
3. We find that antagonistic facilitation is stable in highly stressful conditions, which supports the theory of ecosystem engineers as drivers of primary succession and provides a theoretical ground to investigate facilitation mechanistically in the context of the stress gradient hypothesis.
4. Among all potential causes of stress considered in the model, the key environmental parameter driving changes in the interaction between plants is the proportion of the limiting resource available to plants without mining. This finding represents a challenge for empirical studies, which usually measure the resource input or loss in the system as a proxy for stress. We also find that the total root biomass and its spatial allocation through the root system, often used to measure the nature of the interaction between plants, do not predict facilitation reliably.
5. *Synthesis.* Antagonistic facilitation established between an ecosystem engineer nurse plant and neighbor opportunistic individuals can be evolutionarily stable in stressful environments where ecosystem engineers’ self-benefits from mining resources outweigh the competition with opportunistic neighbors. These results align with theories of primary succession and the stress gradient hypothesis as they show that antagonistic facilitation is stable under environmental stress, but it evolves into mutual interference in milder environments. However, using inaccurate parameters to measure facilitation and stress gradients in empirical studies might mask these patterns.

## 1. Introduction

Plant facilitation contributes to increasing and maintaining local biodiversity (Bulleri, Bruno, Silliman, & Stachowicz, 2016; Cavieres, Hernández-Fuentes, Sierra-Almeida, & Kikvidze, 2016; Wright, Wardle, Callaway, & Gaxiola, 2017) and ecosystem services (Bulleri et al., 2018; Losapio et al., 2021; Martinez-Garcia et al., 2023). In plant communities, ecosystem engineers (Hastings et al., 2007; C. G. Jones, Lawton, & Shachak, 1997) are usually shrubs that colonize harsh habitats such as sand dunes (Bai et al., 2018), recent volcanic substrates (Karadimou et al., 2018), deglaciation lands (Kjær, Olsen, & Klanderud, 2018), or unstable stony grounds (Mori, Osono, Cornelissen, Craine, & Uchida, 2017) during the primary autogenic succession (Clements, 1904, 1916; Connell & Slatyer, 1977). To grow in such conditions, plant ecosystem engineers need to evolve specific traits that allow them to colonize hostile environments where most plants cannot survive (Crain & Bertness, 2006). Once the ecosystem engineer has established, opportunistic individuals that could not colonize the landscape by themselves can establish and proliferate (Verdú, Gómez, Valiente-Banuet, & Schöb, 2021). Following this proliferation, both types of individuals might interact via a non-reciprocal cooperative interaction in which the ecosystem engineer increases the fitness of the opportunist (facilitation), and the opportunist reduces the fitness of its benefactor (interference) via competition for limited resources. This type of interaction is known as antagonistic facilitation (Schöb, Prieto, Armas, & Pugnaire, 2014).

Antagonistic facilitation is frequent (Brooker et al., 2008; Soliveres, Smit, & Maestre, 2015) and can determine the evolution of plant communities (Michalet et al., 2011). However, the evolution and stability of antagonistic facilitation pose an evolutionary dilemma, which makes its ubiquity in nature puzzling. Some models have shown that facilitation among plants only stabilizes if ecosystem engineers cluster and engage locally in mutualistic interactions, hence getting a payback via direct reciprocity *sensu* Nowak, 2006 (Kéfi, Van Baalen, Rietkerk, & Loreau, 2008; Travis, Brooker, Clark, & Dytham, 2006). These results imply that plant facilitation is unstable when ecosystem engineers are surrounded by opportunistic neighbors. Other studies conclude that ecosystem engineers must escape antagonistic facilitation by evolving mechanisms to avoid helping their neighbors (Bronstein, 2009; Brooker et al., 2008). Finally, a third group of studies have suggested that facilitation can only persist when the competitive effect of opportunistic plants is so limited and context-dependent that it has no impact on the evolutionary trajectory of ecosystem engineers (Schöb, Callaway, et al., 2014). In this limit, the net effect of the opportunistic plant on the facilitator individual is practically neutral and hence not antagonistic. All these three families of studies question the evolutionary persistence of plant antagonistic facilitation.

Understanding the diversity of mechanisms by which a plant can facilitate neighbors’ success is crucial to determining the evolutionary fate of antagonistic facilitation. Several, independent mechanisms may override resource competition and lead to net positive effects of one plant over another (Hunter & Aarssen, 1988). Facilitation can occur in any situation where plants experience strain and are, therefore, far from the optimal physiological conditions for their development (Liancourt & Dolezal, 2021). A plant can reduce strain for its own benefit and the benefit of neighbors by driving any environmental parameter value toward its physiological optimum. Plants can, for example, engineer their environment to increase the availability of a scarce resource, such as water or nutrients (Lozano, Armas, Hortal, Casanoves, & Pugnaire, 2017; Francisco I. Pugnaire, Armas, & Valladares, 2004). They can also perform microclimatic ameliorations in many different ways. For example, buffering extreme temperatures (Sánchez-Gómez, Valladares, & Zavala, 2006), buffering atmospheric water demand (Soliveres et al., 2011), and reducing excessive sun radiation through shade (Valladares, Laanisto, Niinemets, & Zavala, 2016). Plants can also protect other individuals growing within their canopies from physical damage caused by abrasion (Okin & Gillette, 2001; Smith, Germino, Hancock, & Johnson, 2003), pests, and diseases (Brooker, Karley, Newton, Pakeman, & Schöb, 2016), and herbivory (Graff, Aguiar, & Chaneton, 2007). Each facilitation mechanism requires separate attention, and we here focus on facilitation via soil resource engineering as a case study.

The root exudation of organic acids, chemical signals, and enzymes can improve soil quality in several ways. Root exudations can dissolve unavailable soil nutrients such as phosphorus (P), iron (Fe), or potassium (K), avoid the toxic effects of hypoxia or an excess of aluminum (Al), and allow the establishment of symbiotic relations with nitrogen (N)-fixing microorganisms (Dakora & Phillips, 2002; Hinsinger et al., 2011; D. L. Jones, 1998). Because N and P are essential resources for plants and are typically scarce in soils, plants that can boost the availability of these macronutrients have received considerable attention (Koffel, Boudsocq, Loeuille, & Daufresne, 2018; Lambers, Raven, Shaver, & Smith, 2008). Plants adapted to grow in P-deficient soils can develop expensive structures, known as cluster roots, that allow them to release the P that has been sorbed to soil particles and make it available for locally foraging roots (Britto Costa, Staudinger, Veneklaas, Oliveira, & Lambers, 2021; Raven, Lambers, Smith, & Westoby, 2018). Some plants growing in N-limited soils can also develop root nodules and establish mutualistic relations with N-fixing microorganisms that trade N for photosynthates (Sprent, 1989; van Velzen, Doyle, & Geurts, 2019). Moreover, some plants can increase soil moisture by having root systems with specific characteristics. For example, plants with roots able to break superficial soil compaction can increase local water infiltration to the soil (Bromley, Brouwer, Barker, Gaze, & Valentin, 1997; Montaña, 1992), and plants with deep tap roots reaching the water table can increase water availability in superficial soil layers by hydraulic lift (Caldwell & Richards, 1989; Prieto, Armas, & Pugnaire, 2012; Zapater et al., 2011). In all these situations, plant ecosystem engineers have specific root traits that allow them to locally ameliorate soil conditions at a cost to themselves, creating islands of fertility where opportunistic plants can potentially grow.

Determining the environmental conditions and underlying biophysical mechanisms that turn antagonistic facilitation evolutionarily stable is crucial for supporting ecological theories that strongly depend on it, such as the stress gradient hypothesis (SGH). Some ecosystems, such as deserts, Mediterranean shrublands, or tropical savannas, are more stressful and poorer in resources than others, such as temperate or tropical forests, even when successionally mature. The SGH predicts that facilitation must dominate in more stressful habitats while interference must prevail in mild conditions (Bertness & Callaway, 1994). Because meta-analyses of studies reporting facilitation over stress gradients did not find consistent support for this theory (Maestre, Valladares, & Reynolds, 2005), several authors suggested that stress could be classified as resource-stress and non-resource-stress and that only the former leads to the SGH predictions (He, Bertness, & Altieri, 2013; Maestre, Callaway, Valladares, & Lortie, 2009). Alternatively, the humped-back SGH suggests that plant facilitation maximizes at moderate, rather than highest, levels of stress (Holmgren & Scheffer, 2010; Maestre et al., 2009; Michalet et al., 2006). Predicting how the evolutionary stability of antagonistic facilitation based on resource dynamics changes across environmental gradients is essential to improve our theoretical understanding of the SGH.

On the other hand, validating the SGH experimentally requires controlling for gradients of environmental stress and measuring one plant’s effect on a neighbor’s fitness. Both these tasks require using reliable surrogates measurable in nature, which might be challenging to define. First, the concept of environmental stress is complex (Körner, 2003; Phillips, 1986) and compromises the empirical study of facilitation and the SGH (He & Bertness, 2014; Liancourt, Le Bagousse-Pinguet, Rixen, & Dolezal, 2017; Richard Michalet, Schöb, Lortie, Brooker, & Callaway, 2014). Changes in resource availability can define environmental gradients for plants (e.g., Laliberté, Zemimik, & Turner, 2014), but these must be measured carefully. For example, some studies use the resource input into the soil, such as fertilization experiments or rainfall gradients, as surrogates of resource availability (e.g., Dohn et al., 2013; Wilson & Tilman, 1993). Other studies, however, use the rate of resource loss, such as nutrient leaching or soil drying rates (e.g., Cabal & Rubenstein, 2018; Pugnaire & Luque, 2001). Both resource input and loss rates determine resource dynamics and can explain its scarcity, but they represent different mechanisms and hence may lead to different plant foraging behaviors (Cabal, Martínez-García, De Castro, Valladares, & Pacala, 2021). Second, given the elusive and non-tangible nature of fitness (Costa, 2013; Primack & Hyesoon Kang, 1989), empirical studies need to rely on indirect measures to quantify how a focal plant influences the fitness of a neighbor. The most common fitness surrogates used by plant biologists are survival rates, reproductive allocation, or biomass production, to name the most common. In particular, many studies consider that biomass production is a good proxy for fitness (Younginger, Sirová, Cruzan, & Ballhorn, 2017) and compare the total biomass (e.g., Dohn et al., 2013) or the spatial distribution of root biomass (e.g., Britto Costa et al., 2021) growing alone vs. interacting to quantify fitness consequences of plant interactions. These studies assume that more biomass or allocation of biomass toward the neighbor when interacting as compared to alone indicates that plants are being facilitated. A mechanistic understanding of antagonistic facilitation based on resource dynamics would help to determine which of the parameters that can be measured in the field or controlled experimentally are better to identify facilitation and define stress gradients.

Empirical studies reporting facilitation among plants in specific systems are copious, but we still lack a solid theoretical ground establishing general principles (Zhang & Tielbörger, 2020). In particular, we lack mechanistic models that unravel the biophysical processes sustaining net positive interactions among plant individuals and how those interactions change across environmental gradients (Brooker et al., 2008; Soliveres et al., 2015). Here, we develop such a mechanistic model for the specific case of individual plants interacting via a shared soil resource in explicit space. Following this modeling approach, plant interactions are an emergent property of resource use rather than postulated a priori via parameter choices. We use this model to answer three questions. First, can ecosystem engineers establish evolutionarily stable antagonistic facilitation interactions with opportunistic plants competing for the same resource? Second, is antagonistic facilitation evolutionarily stable in highly or moderately stressful environments but disappears in mild habitats? If so, what are the key biophysical mechanisms underlying a soil resource-driven SGH? Finally, according to our model results, which environmental parameters and fitness surrogates are better for the empirical study of facilitation and the SGH?

## 2. Materials and Methods

### 2.1 Plain text summary of the model

We model the below-ground root growth of plant individuals sharing neighborhood areas and competing for a single limiting resource. We assume that the total resource input splits into two pools: unavailable and available resource, and consider two types of plants: soil ecosystem engineers and opportunistic individuals. Both types of plants forage from the available resource pool (e.g., water that infiltrates in the soil), and ecosystem engineers can mine from the unavailable pool and render resources available for all plants (**Figure 1a**). We further assume that the fraction of unavailable resource (e.g., runoff water) is lost if not mined instantaneously and therefore does not accumulate over time (e.g., plant-triggered increase in infiltration). The available resource also leaves the soil at a constant rate due to physical losses (e.g., percolation, evaporation). Given this resource dynamics, plants spread their root system (root biomass per volume unit of soil) to maximize their net resource gain by balancing the benefits obtained from the resources they can forage and the root production costs. The amount of resource plants can forage depends on the input and physical loss of available resource, total mining intensity, and foraging by other plants. Root production costs increase with the distance to the plant stem for any type of plant (representing the need to grow longer and thicker transportation roots) and, for ecosystem engineers, with mining intensity.

**Figure 1:**
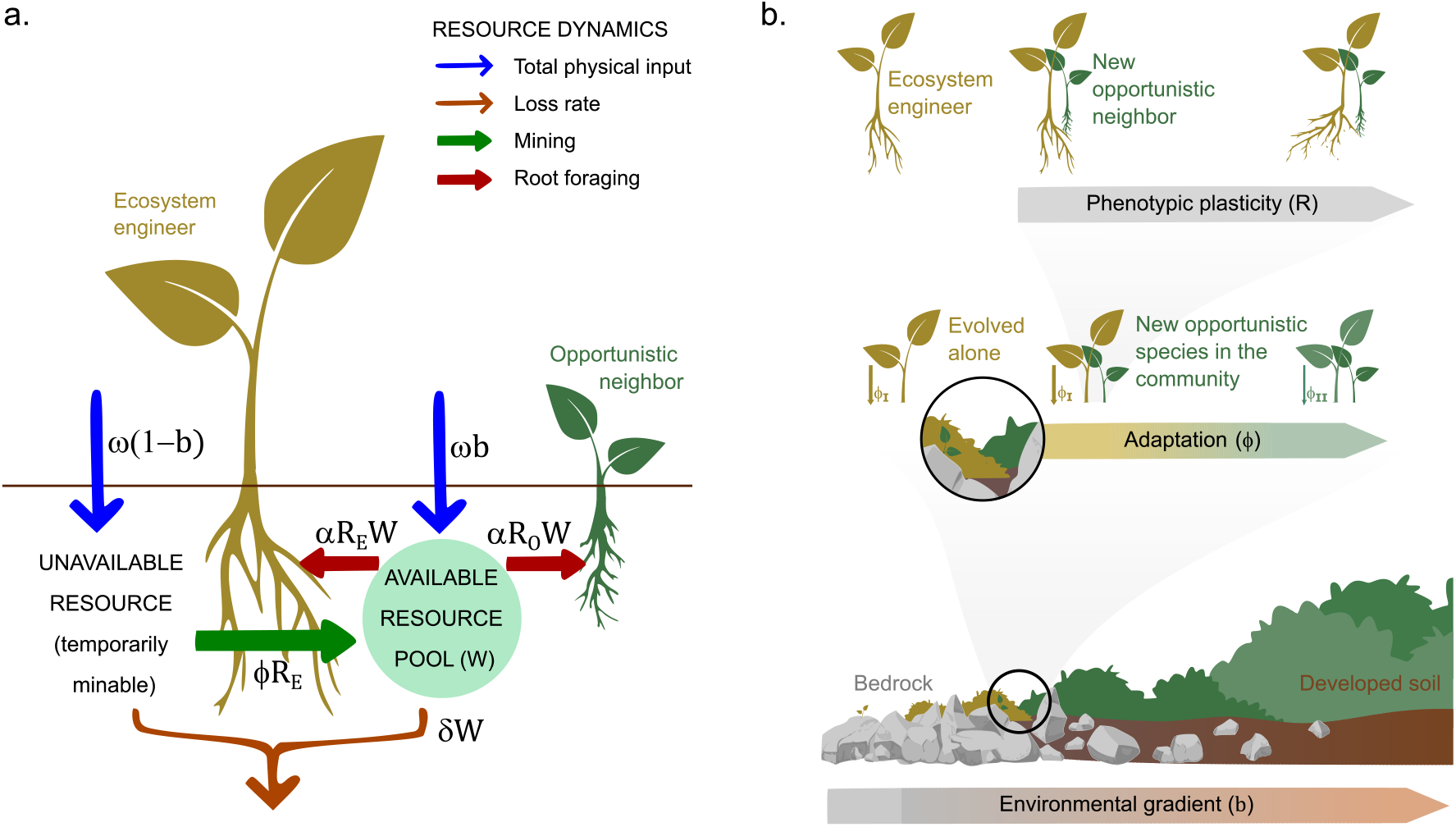
**a.** Schematic representation of soil resource dynamics in a specific soil location in our model. This dynamics is replicated at each point of space labeled with ℓ in the model equations. We consider a constant input of potential resource (*ω*) that splits into spontaneously available to plants (*b*) and temporarily minable (1-*b*). Ecosystem engineer roots can mine the latter and make it available to all plants at a rate proportional to the mining intensity (*ϕ*) and the ecosystem engineer root density (*R*_E_). Every plant can forage the available resource *W* (in quasi-equilibrium) at a rate proportional to their per-capita absorption rate (*α*) and root density (*R*_E_ for the ecosystem engineer and *R*_O_ for the opportunistic). The available resource also leaves the soil at a physical decay rate (*δ*). **b.** The different scales at which we assessed the emergence of facilitative interactions, including the plastic response of plant root distribution *R* at the life span time scale (top), the adaptive response of the ecosystem engineers’ facilitator trait *ϕ* at the evolutionary time scale (center), and the change in the fraction of soil resource that is spontaneously available over succession *b*, which can change through space or at a geological time scale (bottom).

Complete solutions of this model provide both the optimal root distribution for single or multiple plants and, for ecosystem engineers, the optimal value of the resource mining trait. We calculate both quantities at equilibrium, when plants are fully grown and producing additional roots would decrease their net resource gain. We first obtain the optimal spatial distribution of root density that maximizes the net resource gain at each soil location. Next, we use this root density distribution to calculate the net resource gain for a plant at each point of soil. Third, integrating the spatial distribution of root density and the net resource gain across the extension of the root system, we calculate the plant-level root biomass and plant-level resource net gain, respectively. Lastly, we calculate the optimal mining intensity as the trait value that maximizes the plant-level net resource gain for the root density distribution obtained in the first step.

We can obtain these model outputs for different setups and parameterizations. For example, we run the model using both a single plant and a pair of plants and compare the plant-level net resource gains of a chosen focal plant in these two scenarios. Based on this comparison between plant-level net resource gains in different growing environments, we determine whether plants interact via facilitation (the plant-level net resource gain of the focal plant is higher when it grows paired than when it grows alone) or interference (otherwise). Additionally, for the ecosystem engineers, we assess the evolution of the mining trait under different environmental conditions and for engineer plants growing alone or in the presence of opportunistic plants. Lastly, we explore how root spatial distributions and optimal mining intensities vary with the different environmental parameters: total resource input, proportion of such resource that is available spontaneously without mining, and physical rate at which the resource leaves the soil. Different environmental parameterizations represent different scenarios that can be interpreted as stress gradients in space or across successional time. Therefore, through its various variables and parameters, our model accounts for processes occurring at different time scales: plants’ phenotypically plastic root distribution, the evolution of the ecosystem engineer trait, and changes in the abiotic conditions across an environmental gradient (that can be defined in space or time) (**Figure 1b**).

### 2.2 Mathematical model formulation

We extend the model for resource competition introduced in Cabal, Martínez-García, De Castro Aguilar, Valladares, & Pacala, 2020 to investigate the evolution of plant antagonistic facilitation. Because we model individual root growth and spatial allocation, we use a spatially explicit modeling approach. The model assumes a series of processes governing the dynamics of a soil resource *W* and provides the net resource gain for each plant due to the balance between how much resources it uptakes at a soil location, modeled via a classical consumer-resource interaction term, and the cost of foraging in that soil patch. It therefore consists of a series of consumer-resource models at each soil patch, with roots from different plants competing for a single limiting resource. We consider two types of plants: Opportunistic plants, that can only forage soil resources, and plant ecosystem engineers that consume soil resource and also its availability by evolving a resource mining trait.

#### Resource dynamics

We model resource dynamics in each soil point as a combination of three processes: input at rate *I*, abiotic loss at rate *δ*, and resource uptake by foraging roots *R* at a per capita rate *α* (see **Table 1** for a summary of the environmental parameters and their dimensions). For mathematical simplicity, we consider a linear, non-saturating resource uptake term and that both ecosystem engineer and opportunistic plants have the same per capita uptake rate. More complex scenarios, which are not the focus of this study, could account for plant type-specific per capita uptake rates and nonlinear, saturating uptake terms, including tradeoffs in resource uptakes between plant types (Grover, 1991; Tilman, 1982; Vincent, Scheel, & Brown, 1996). To incorporate the effect of ecosystem engineers, we model the resource input as a combination of spontaneously available resource and unavailable resource that is temporarily available for mining. In soils lacking ecosystem engineering roots, only a fraction *b* of all the potential resource, *ω,* is spontaneously available to roots (0 ≤ *b* ≤ 1). Ecosystem engineer roots *R_E_* increase the fraction of *ω* that is available for plant foraging due to a per capita mining intensity *ϕ*. We model this positive feedback as

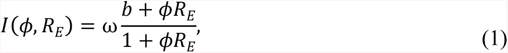

which saturates at *ω* when the mining intensity tends to infinity to account for the finiteness of resources. Putting all these terms together, the resource concentration changes at each soil location according to

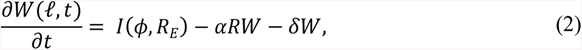

where we have dropped the dependence on space and time from the terms on the right side of Eq. (2) and *ℓ* is the spatial coordinate of the soil location measured from the insertion point of the focal plant’s stem to the soil surface.

**Table 1:** Parameter description and values (if not indicated otherwise) for the environmental resource dynamics.

| Parameter (Environment) | Symbol | Units | Value |
| --- | --- | --- | --- |
| Potential resource input | $\omega$ | ml.day<sup>-1</sup> | 5 |
| Proportion of $\omega$ abiotically available | b | - | 0.05 |
| Resource abiotic decay rate | $\delta$ | day<sup>-1</sup> | 0.1 |

#### Net resource gain

At each soil location, the net resource gain of an individual plant, *j*, at time *t* is the balance between the resource uptake and the cost associated with growing and maintaining the roots needed to forage in that soil location,

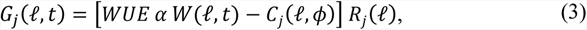

where *R* is the foraging root density (units of fine root biomass per unit of soil volume), *C* is the cost of producing and maintaining such root density, and *WUE* is the resource use efficiency that represents the conversion factor from harvested resource to new root biomass.

Because the cost function depends on the distance between the soil patch and the plant insertion point to the soil surface, the model is spatially explicit. We account for three contributions to the cost function: the cost of producing and maintaining fine roots that absorb resources from the soil *c_r_*, the cost of building and maintaining transportation roots that connect the fine roots to the plant stem *c_t_*, and the cost per density of fine roots associated to a unit increase in the mining intensity *c_e_*,

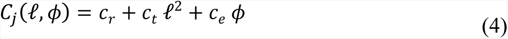

For opportunistic plants, *ϕ* = 0 and the cost function in Eq. (4) has only two contributions (see **Table 2** for a summary of the plant parameters) (Cabal et al., 2020).

**Table 2:** Parameter description and values for the plant functional traits in spreading opportunistic plants (SPR), normal opportunistic plants (NOP), and ecosystem engineer plants (EEP).

| Parameter (Functional traits) | Symbol | Units | SOP | NOP | EEP |
| --- | --- | --- | --- | --- | --- |
| Uptake rate per unit fine root weight | $\alpha$ | ml.mg<sup>-1</sup>.day<sup>-1</sup> | 10 | 10 | 10 |
| Resource use efficiency | WUE | ml.mg<sup>-1</sup> | 1 | 1 | 1 |
| Cost of fine roots | $c_r$ | ml.cm<sup>3</sup>.mg<sup>-1</sup>.day<sup>-1</sup> | 5 | 5 | 5 |
| Cost of transportation roots | $c_t$ | ml.cm<sup>3</sup>.mg<sup>-1</sup>.day<sup>-1</sup> | 0 | 0.2 | 0.2 |
| Cost of facilitation trait | $c_e$ | ml.cm<sup>3</sup>.mg<sup>-1</sup>.day<sup>-1</sup> | 0 | 0 | 1.25 |
| Resource-mining trait | $\phi$ | ml.mg<sup>-1</sup>.day<sup>-1</sup> | 0 | 0 | $\phi_I, \phi_{II}$ |

### 2.3 Two-step model analysis

Three parameters determine soil resource physical dynamics in our model and therefore define the environmental stress: the potential resource input *ω*, the fraction of such resource that is spontaneously available to plant roots *b*, and the resource physical loss rate *δ*. For example, if we consider that water is the limiting resource, *ω* could represent precipitation at a given location; *b* and *1-b*, the fraction of the water input that infiltrates in bare soil and is lost by runoff, respectively; and *δ,* the loss rate due to percolation to the subterranean water table where roots do not reach. Similarly, if *W* represents a soil nutrient, *ω* could account for the amount of such nutrient present in the rock or mineral particles interface; *b* and *1-b,* the proportion of this nutrient that spontaneously dissolves in pure water and remains sorbed and the chemically unavailable proportion, respectively; and *δ,* the rate at which this nutrient is leached with water to the subterranean water table and leaves the soil system. We can evaluate the outcome of the pairwise plant interaction across environmental gradients defined by changing values of *ω, b,* and *δ,* and thus representing each of these three different mechanisms.

We conducted a two-step computational analysis for different values of these parameters. In the first step, we obtained the resource mining trait value that optimizes the plant-level net resource gain and is thus expected to have evolved in those environmental conditions. We used this result to parameterize the model using a resource mining intensity expected to evolve in the environmental conditions defined by the values *ω*, *b*, and *δ*, and perform the second step of the analysis. In this second step, we quantified the plant-plant interaction simulating experiments at the plant life span timescale using the resource mining intensity obtained previously. We performed this two-step analysis both for solitary and interacting plants.

To perform all the computational analysis, because the resource dynamics is much faster than root growth, we assumed that the resource concentration is in quasi-equilibrium in each step of plant growth. This approximation allows us to write a closed expression for the net resource gain of each plant at a given soil location (see SM for a complete derivation),

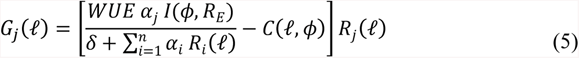

where *R_E_* is the total root biomass density of the ecosystem engineer at location ℓ. To optimize *G* in a given environmental scenario, we look for combinations of root density profiles and resource mining intensity that optimize the net resource gain in Eq. (5).

#### 2.3.1 First step: model parameterization

To mimic natural conditions in which ecosystem engineer plants might have evolved the resource mining trait, we first allowed*ϕ* to evolve either assuming that ecosystem engineers grew alone, *ϕ_I_,* or competing with opportunistic plants, *ϕ_II_*. We obtained these values of *ϕ* for a range of environmental stresses, e.g., from *b* = 0 (high stress) to *b* = 1 (low stress). This first analysis yields the evolutionarily stable values of the resource mining trait for an ecosystem engineer plant at a given environmental stress level and assuming that the ecosystem engineer adapted to growing alone or surrounded by opportunistic plants.

Modeling a community considering several individual opportunistic and ecosystem engineer plants in interaction with one another would make the model mathematically intractable and require many arbitrary choices about community size and spatial structure. Instead, to get a general competitive background for the evolution of *ϕ_II_*, we modeled the opportunistic individual as a spreading plant that grows vegetatively covering the soil surface (de Kroons & Hutchings, 1995). These spreading opportunistic plants distribute their stem everywhere on the soil surface, like a stolon or a rhizome, so their roots grow across the canopy area of the ecosystem engineer without any spreading costs (Dolezal et al., 2019). Mathematically, this growth strategy implies removing the costs associated with transportation roots, *c_t_* = 0. We consider this assumption a more realistic and tractable representation of the average background community to which ecosystem engineers may have adapted. We provide a detailed description of how this analysis was performed in the Supplementary Material.

#### 2.3.2 Second step: simulated experiments

In a second step, we considered individual plants from the above-parameterized conditions, i.e., using the values *ϕ_I_* or *ϕ_II_* expected to evolve at given environmental conditions, and simulated experiments with such plants and at such environmental conditions at the scale of their life span. This analysis measures the nature of the pairwise plant interaction. We performed two classes of independent experiments. First, we simulated the growth of an opportunistic plant alone. Second, we simulated the growth of an opportunistic-engineer pair of interacting plants separated by a distance *d*. To maximize the strength and effects of the interaction, we mostly assumed that both interacting plants shared the same insertion point to the soil (*d* = 0), but we also measured the outcome of the plant interaction for *d*=5cm (**Fig. SM3**). We obtained the root biomass distribution *R(ℓ) for both solitary and paired opportunistic plants*. We calculated the plant-level net resource gain associated with such root distribution, 𝒢, integrating *G* [Eq. (5)] in space. Finally, using the results from the simulated experiments considering solitary and interacting plants, we estimated the net interaction from the ecosystem engineer to the opportunistic plant by comparing the plant-level net resource gain of solitary and paired opportunistic individuals. Following a similar procedure, we also calculated the total root biomass of plants, ℜ, by integrating the local *R(ℓ)* values across space and estimated the difference in root proliferation by comparing the values for solitary and paired plants. To quantify both the net interaction from the ecosystem engineer to the opportunistic plant and to which extent the opportunistic plant changes its root biomass in the presence of the ecosystem engineer, we calculated coefficients based on the normalized ratio *RII* (Armas, Ordiales, & Pugnaire, 2004):

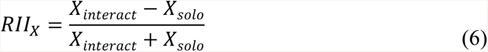

where *X* is ℜ or 𝒢 depending on whether we measure changes in plant-level root biomass or net resource gain, respectively. *X_interact_* is the value obtained for ℜ or 𝒢 when the opportunistic plant grows close to an ecosystem engineer and *X_solo_* is the value obtained for solitary plants. *RII* can vary between −1 and 1. *RII*_𝒢_ informs about the net interaction, with *RII*_𝒢_ = −1 indicating that the plant cannot coexist with the neighbor (competitive exclusion); −1 < *RII*_𝒢_ < 0 net interference; *RII*_𝒢_ = 0 no effect; 0 < *RII*_𝒢_ < 1, net facilitation; and *RII*_𝒢_ = 1 indicating that the plant cannot exist without the neighbor (obligatory facilitation). For root biomass measurements, *RII*_ℜ_ < 0 (*RII*_ℜ_ > 0) indicates that opportunistic plants under-proliferate (over-proliferate) roots when interacting with an ecosystem engineer relative to plants growing alone.

We hypothesize that the net plant interaction changes more substantially over an environmental gradient defined by changes in *b* because this parameter controls how much of the total resource input becomes readily available for plants and how much goes to the temporarily mineable resource pool and is accessible only to soil engineer roots (Fig. 1a). Hence, *b-*stress causes a strain that can be reduced by soil engineer roots. On the other hand, *ω* and *δ* determine the total input and loss rate, and create a strain that ecosystem-engineer roots cannot mitigate. Therefore, we first analyze how the net plant interaction changes across an environmental gradient defined by changes in *b* and then conduct a more extensive scanning of the three-dimensional parameter space also to quantify the role of *ω* and *δ* in shaping the pairwise plant interaction.

## 3. Results

### 3.1 Evolution of the mining trait and resulting net interactions with changes in b

Only ecosystem engineers with a mining intensity above a given threshold can survive alone in stressful habitats where the proportion of resource that is spontaneously available, *b,* is below a critical value *b_c-o_*. Neither ecosystem engineers investing in resource mining below that threshold nor opportunistic plants, for which the mining trait *ϕ=0*, can survive because their root production and maintenance costs outweigh resource uptake. Opportunistic plants can invade the ecosystem in this regime only at the shelter of ecosystem engineers, hence establishing an obligatory interaction with them (**Figure 2, I**). Above the critical value *b_c-o_* opportunistic plants can invade the environment without interacting with the ecosystem engineers (**Figure 2, II** with a hypothetical invasion event represented by the red dot). The optimal resource-mining intensity for solitary ecosystem engineers is a finite value *ϕ_I_* that depends on the environmental stress defined by the value of *b*. Mining intensities greater than *ϕ_I_* are suboptimal because the costs of increasing the mining capacity outweigh the revenue per increased unit. Indeed, at very high values of *ϕ*, the plant-level net resource gain and the optimal root biomass become zero because the cost of producing roots with such mining ability exceeds the amount of resource they can mine and uptake, and the plant cannot establish. The optimal value of the resource mining trait for solitary individuals, however, does not coincide with the value of *ϕ* at which root biomass is maximal. Values of *ϕ* lower than *ϕ_I_* result in more roots but with lower plant-level net resource gain (**Figure SM1**).

**Figure 2:**
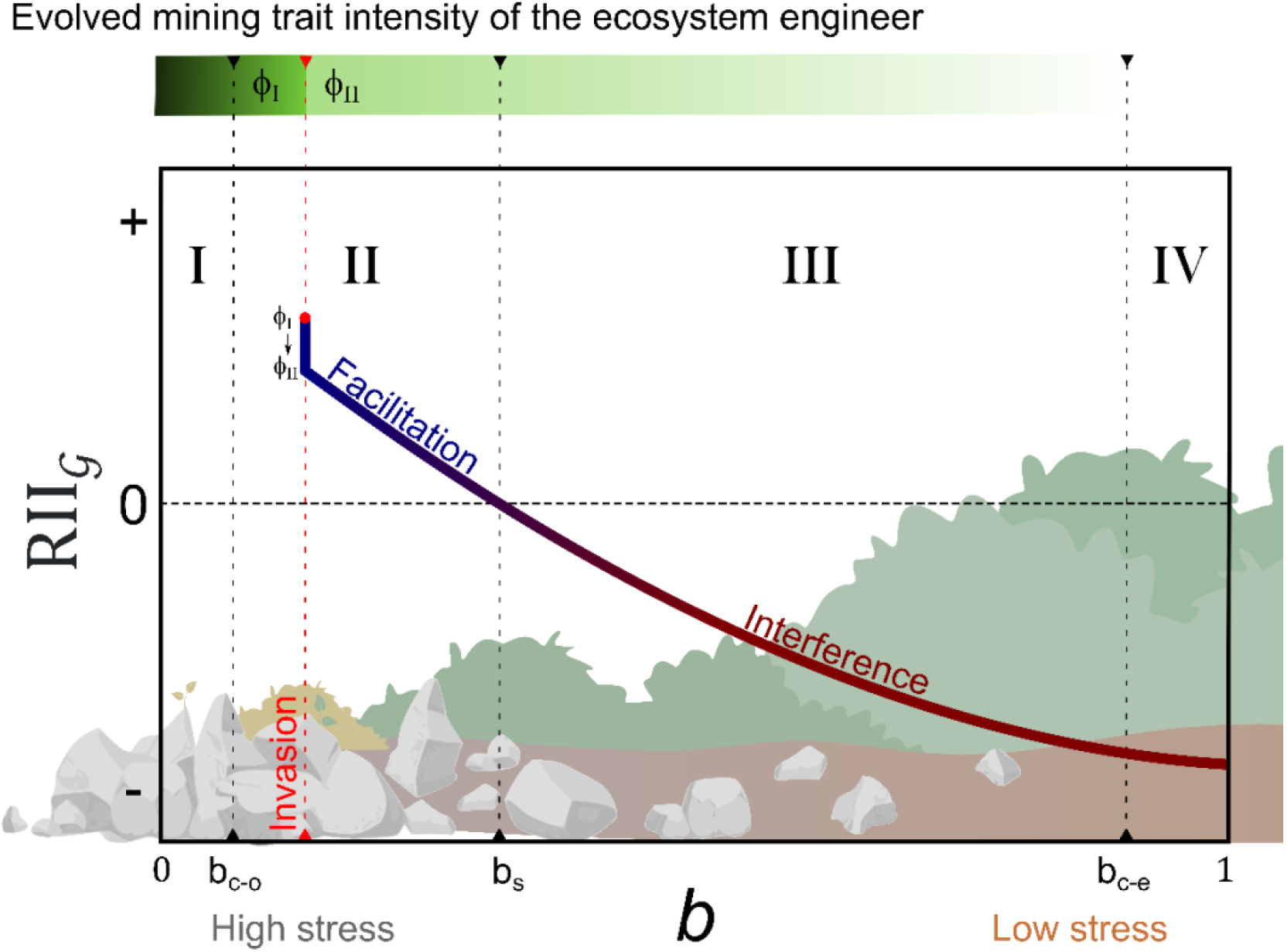
Graphical summary of the changes across an environmental gradient defined by the proportion of resource that is spontaneously available to plant roots, *b* (see **Figure SM2** for detailed results obtained numerically from model equations). Across the gradient, *b* varies from 0 (high stress, none of the resource is available without mining) to 1 (low stress, all the resource is spontaneously available). The figure shows the evolution of the mining trait *ϕ* in the ecosystem engineer (top bar, darker green represents higher values of *ϕ*) and the net interaction of the ecosystem engineer on the opportunistic plant measured using the normalized ratio of net gain for the opportunistic plant *RII*_𝒢_ (central figure). Across the environmental gradient, we distinguish several regions. (I) In highly stressed environments, ecosystem engineers colonize the ecosystem and evolve an intense mining trait *ϕ_I_*, which progressively decreases as *b* increases and environmental conditions improve. (II) Passed *b=b_c-o_*, opportunistic plants do not need to interact with ecosystem engineers to invade the system. Following the invasion of an opportunistic plant (shown in the figure by a red dot at an arbitrarily chosen value of *b* in this region, where we consider it most likely to occur), ecosystem engineers evolve to adapt to the presence of opportunistic neighbors and the mining trait intensity suddenly drops from *ϕ_I_* to *ϕ_II_* (sharp transition in green darkness in the top bar). However, *ϕ_II_* remains positive and soil resource engineering persists. (III) As environmental conditions improve further and *b* surpasses *b_s_*, antagonistic facilitation disappears and all plants interfere with each other. In this regime, there is no nursing effect even though *ϕ_II_* remains positive, and soil resource engineering persists. (IV) Finally, *ϕ_II_* vanishes when *b>b_c-e_*, indicating that ecosystem engineers stop existing (mining evolves to disappear) and all plants act as opportunistic individuals that compete for resources.

**Figure 3:**
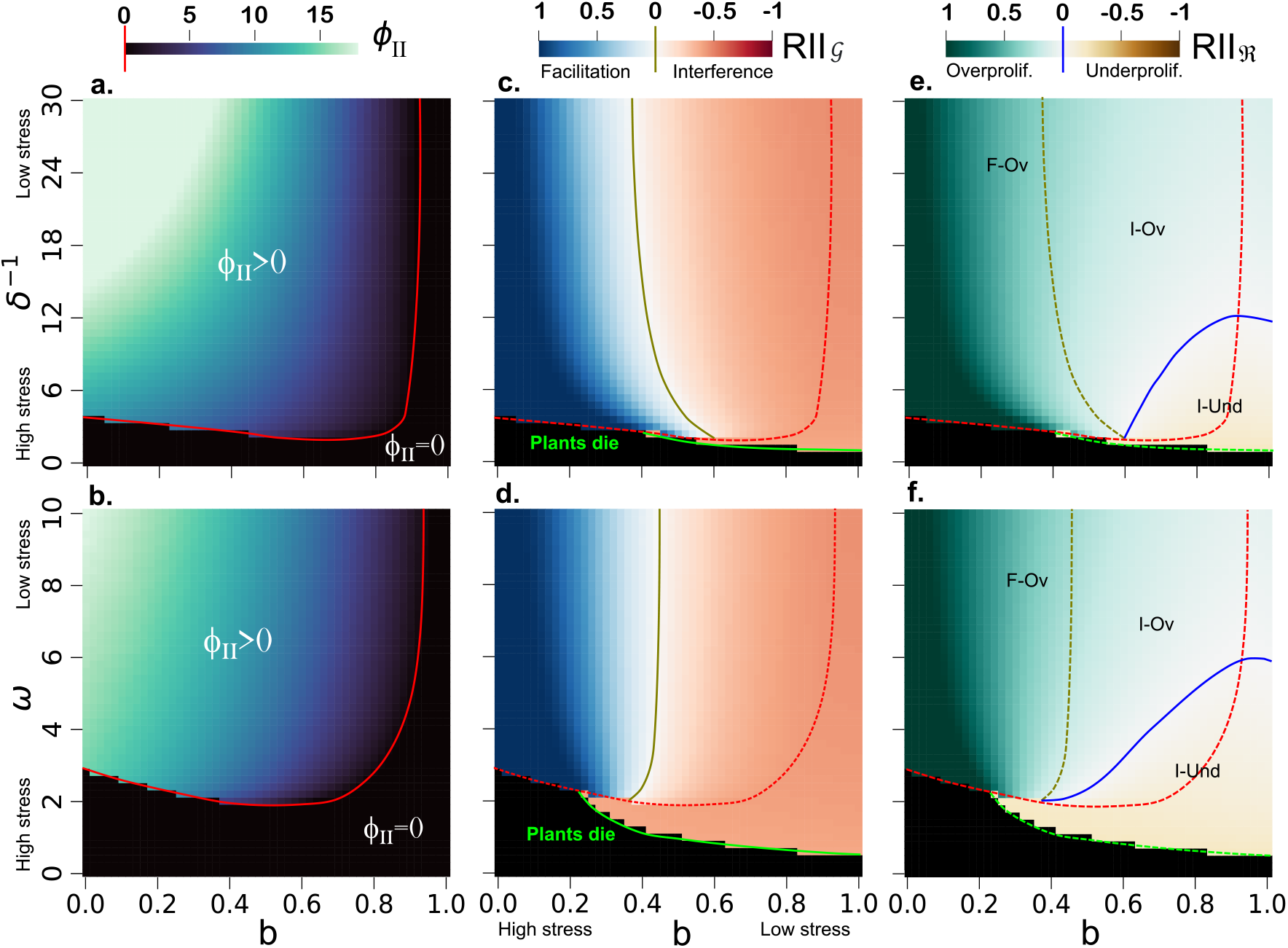
For an interacting pair of ecosystem engineer/opportunistic plants, changes in the resource-mining trait *ϕ_II_* of the ecosystem engineer (**a, b**); RII for the opportunistic plant-level net resource gain *RII*_𝒢_ (**c, d**); and root biomass *RII*_ℜ_ (**e, f**). We measure each of these changes at different values of the three environmental parameters determining the physical dynamics of the solid resource: the proportion of available resource *b*, total resource input *ω* and physical resource loss *δ*. The letters in the rightmost panels indicate regions of the parameter space with Facilitation (F) or interference (I) and with root Overproliferation (Ov) or underproliferation (Und).

If opportunistic plants invade these stressful habitats (red dot in Fig. 2), ecosystem engineers transiently have a strong positive effect on their opportunistic neighbors because they maintain the resource mining intensity that evolved in the absence of opportunistic plants (*ϕ_I_*). However, opportunistic plants change the resource dynamics due to competition for the shared resource. These new processes driving resource dynamics impose a strong evolutionary pressure on the resource mining trait and trigger a post-colonization shift in the optimal trait value from *ϕ_I_* to a lower value *ϕ_II_*. This change in the evolved value of the resource mining trait (*ϕ_I_* →*ϕ_II_*) increases the ecosystem engineer’s net gain and decreases the interacting opportunistic plants’ net gain (**Figure SM2**). Once the ecosystem engineer has adapted to the presence of opportunistic neighbors and evolved a new value for the mining trait, antagonistic facilitation is evolutionarily stable for any *b* below a critical shifting point *b_s_* (**Figure 2, II**). However, individuals able to mine resources keep a non-zero *ϕ_II_* value indicating that they still act as ecosystem engineers. If opportunistic plants invade extremely stressed environments with *b* < *b_c-o_*, ecosystem engineers allow their survival within their neighborhood area. If they invade when *b_c-o_* < *b* < *b_s_*, ecosystem engineers will increase the opportunistic plants’ net gain compared to their performance in bare soil. Therefore, ecosystem engineers also act as nurse plants in both cases, and antagonistic facilitation is evolutionarily stable (*RII*_𝒢_ > 0).

Above *b_s_*, ecosystem engineers keep their mining trait *ϕ_II_* positive, meaning that they are still acting as ecosystem engineers, but the plant-level net resource gain of an opportunistic plant is higher when it grows alone than near an ecosystem engineer (*RII*_𝒢_ < 0; **Figure SM2**). Hence, the ecosystem engineer is no longer a nurse plant. In this case, the strength of exploitative competition by ecosystem engineers exceeds the positive effect of the mining trait on the opportunistic plant and the interaction becomes mutual interference (**Figure 2, III**).

As 𝑏 increases and more resource is spontaneously available for plants, the resource that ecosystem engineers can mine becomes progressively lower and *ϕ_II_* decreases. This decrease in the optimal mining intensity also reduces the difference between the plant-level net resource gain of solitary ecosystem engineer and solitary opportunistic plants (**Figure SM2**). When *b* reaches a critical value *b_c-e_*, the optimal mining intensity becomes *ϕ_II_* = 0, indicating that ecosystem engineers evolve to lose the resource mining trait and become opportunistic (**Figure 2, IV**).

### 3.2 The different effects of environmental parameters on plant interactions and root proliferation

Through the three-dimensional parameter space defined by *b*, the physical loss of resource *δ*, and the potential resource input *ω* and encompassing all possible environmental conditions, we can identify a number of regions in which plant survival and the biotic interaction vary qualitatively. For fixed values of *δ* and *ω*, *ϕ_II_* in ecosystem engineer plants decreases progressively as *b* increases (stress decreases), until *ϕ_II_* = 0 at low stress (*b* > *b_c-e_*). However, *ϕ_II_* changes abruptly from 0 at high levels of *δ*- and *ω*-stress, to a positive value *ϕ_II_ >* 0 when stress levels in both parameters go below a threshold (**Fig 4a, b**).

The *RII* coefficient for the plant-level net resource gain of opportunistic plants *RII*_𝒢_ shows that the interaction transitions gradually from facilitation at low values of *b* (high *b*-stress) to interference at higher values of *b*. In contrast, at very high *δ*- and *ω*-stresses not even engineers can survive (**Fig 4c, d**) because the stress is induced by resource loss that cannot be mined. As the stress induced by these two parameters decrease, there is a first environmental threshold (solid green lines in **Fig. 4c, d**) above which ecosystem engineers can survive but do not evolve a positive value in their mining trait. In that range of the environmental gradient, defined by moderately high levels of *δ*-stress and *ω*-stress, pairs of exploitative plants (because ecosystem engineers behave as purely exploitative) compete similarly to what we observe for *b* ≥ *b_c-e_*. At even lower *δ*- and *ω*-stress we find a second threshold where ecosystem engineers acquire their positive mining trait in an abrupt shift (dashed red lines in **Fig. 4c, d**). After passing that second threshold, the net interaction, measured by *RII*_𝒢_, depends strongly on *b* but barely changes with *δ* and *ω*. Only for a very narrow range of intermediate values of *b*, the interaction changes slightly from facilitation to competition as *δ*-stress decreases, but from competition to facilitation as *ω*-stress decreases. We observe that the fraction of resource that is available without the intervention of mining, 𝑏, rather than the potential resource input *ω* or the physical decay rate of the resource 𝛿, accounts for most of the variation in the sign and the strength of the biotic interaction in the environmental region when pairs of ecosystem engineer/opportunistic plants interact.

The analysis of root production using *RII_R_* shows that the transition from facilitation to competition (dashed golden lines in **Fig. 4e, f**) does not coincide with the transition from root overproliferation to underproliferation (solid blue line in **Fig 4e, f**). This result allows for three possible interaction outcomes. First, when there is antagonistic facilitation the opportunistic plant always overproliferates roots (F-Ov). Second, plants might interact via mutual interference, but the opportunistic plant still overproliferate roots as compared to growing alone (I-Ov). Finally, plants might interfere with each other and underproliferate roots (I-Und). The latter two interaction outcomes can happen for positive values of *ϕ_II_* (active mining of the ecosystem engineer) or for *ϕ_II_* = 0 (both plants behave as purely exploitative, i.e., opportunistic). In addition to that, we found that a skewed distribution of the opportunistic plant’s root density in space toward the ecosystem engineer can also be observed in both scenarios, facilitation and interference (**Fig SM3**).

## 4. Discussion

### 4.1 Successional changes in plant biotic interactions

For facilitation via soil resource engineering, the amount of resource available to plants without the intervention of root mining is the leading force driving the change in plant interactions across environmental gradients. Only ecosystem engineers can colonize highly stressed ecosystems, which could represent the initial stages of succession at which opportunistic plants cannot survive on their own. Nevertheless, after ecosystem engineers colonize the system, opportunistic plants can establish an obligatory interaction with their benefactors and invade the ecosystem. Previous work has suggested that ecosystem engineers may lose their mining trait over the course of evolution due to the presence of opportunistic neighbors that benefit from mined resources at no cost (Koffel et al., 2018; Song, Yu, Jiang, Korpelainen, & Li, 2019; Walker & Chapin III, 1987). Our results, however, suggest that antagonistic facilitation is evolutionarily stable at high environmental stresses. In this regime, ecosystem engineers benefit from bearing the mining trait despite the presence of opportunistic plants and they have an overall positive effect on opportunistic neighbors because mining benefits overcome resource competition. In milder environments, ecosystem engineers still mine resources from unavailable pools for their own benefit, but resource competition dominates over mining benefits and the net interaction between the ecosystem engineer and the opportunistic plant transitions from antagonistic facilitation to mutual interference. In this regime, ecosystem engineers are not ‘nurse plants’ and become pure competitors for opportunistic plants. Finally, at very low environmental stress mimicking the conditions of well-developed soils where most of the resource is spontaneously available to the plants, ecosystem engineers evolutionarily lose their mining trait, and all plants evolve an opportunistic strategy.

### 4.2 Consequences for the stress gradient hypothesis

Our model further dissects stress induced by resource availability into three parameters; the total resource input *ω* (e.g., levels of precipitation), the fraction of such resource that becomes available to plants spontaneously *b* (e.g., how much of the rainwater infiltrates in bare soil and becomes available to roots without the intervention of ecosystem engineers), and the rate of physical resource decay *δ* (e.g., how fast the infiltrated water leaves the soil due to evaporation or percolation). Facilitation can occur only if there is strain, and ecosystem engineers can reduce such strain. Contrarily, if the physical environment is near optimal, or if plants cannot improve its quality, facilitation is not possible. Our results support the SGH for resource stress if the stress is caused by the fraction of the total resource that is available for plants without mining *b*, but we did not find a significant impact of the other environmental parameters, the physical decay rate *δ* and total resource input *ω*, on the net interaction. This is because, even though all parameters can produce strain, only under *b-*stress resource pools can be mined by ecosystem engineers to reduce strain. As a result, *b* is the main driver of shifts in the biotic interaction between an ecosystem engineer and its opportunistic neighbor across environmental gradients and moderate changes in this parameter would dominate over changes in any other environmental parameter and mask their effects on plant biotic interactions.

Moreover, our results predict that ecosystem engineers abruptly lose their mining trait when *δ* or *ω* control the environmental stress, supporting the humped-back SGH. Within a narrow range of intermediate values of *b*, increasing *δ-* or *ω-*stresses can cause a sudden transition in the plant biotic interaction from interference between two opportunistic plants to facilitation where one plant becomes the ecosystem engineer, followed by a gradual loss of the net facilitation. Teasing apart the contributions of all three parameters to resource dynamics will allow researchers to elaborate more mechanistic descriptions of resource stresses and therefore develop more comprehensive theories regarding the importance of facilitation across a stress gradient.

### 4.3. Consequences for empirical studies

Our results have also important consequences for the interpretation of empirical studies addressing interaction shifts across environmental gradients. We have identified two major sources of potential confusion based on our model. First is the fact that *b* is the main environmental driver of shifts in plant interactions. This parameter, which could represent, for example, the ratio between water infiltration and runoff, is much harder to determine empirically than the other two environmental parameters in our model, *ω* or *δ* (e.g., rainfall or soil drying rates). Many empirical studies assess the change in plant interaction across environmental gradients using the latter two parameters, which our model suggests could lead to misleading conclusions. Second, our calculations of the plant-level net resource gain suggest that plant biomass is a misleading fitness surrogate to determine the net effect of one plant on another. Specifically, root over-proliferation or root density skewed in space towards a neighboring ecosystem engineer might not be a univocal consequence of facilitation, as both behaviors may be observed when interacting negatively with the neighbor due to a tragedy of the commons (Cabal, 2022; Gersani, Brown, O’Brien, Maina, & Abramsky, 2001). This result represents a challenge for experimental designs, because the plant-level net resource gain is not easy to determine empirically. In cases where this measure is not possible, other fitness proxies, such as resource allocation into reproductive tissues, may be better fitness indicators (Kozłowski, 1992; Younginger et al., 2017).

## 5. Conclusions

Antagonistic facilitation enables more diverse plant communities in stressful environments at the initial stages of primary succession and may accelerate the development of mature soils. It also may promote plant biodiversity in harsh habitats, such as deserts or Mediterranean shrublands, where nurse plants facilitate the survival of other species that depend on them. To mechanistically understand the plant-plant interaction and how it shifts across resource-stress gradients, it is crucial to distinguish the biophysical parameters underpinning soil resource availability. Using a mechanistic model of interacting plant individuals we have proved that antagonistic facilitation can be evolutionarily stable in when plants are subject to certain environmental stresses. Moreover, we have shown that, for the specific case of facilitation via soil resource engineering studied here, the main parameter explaining shifts in plant-plant interactions across resource-stress gradients is the proportion of the existing resource that becomes available to roots spontaneously, such as the proportion of rainfall that infiltrates in soil instead of running-off when the resource is water. The amount or resource input (such as rainfall when the resource is water) or the physical decay (such as loses of soil water through evaporation or percolation) play a secondary role. Similarly, the use of biomass production as a proxy of plants’ fitness and hence for measuring plant net interaction is biased because biomass over-δproliferation can result from both facilitation and a tragedy of the commons. Accounting for this level of mechanistic detail in mathematical models for plant interactions and measuring these parameters in field studies is key to better understanding the fundamental drivers of plant-plant interactions.

Our model focuses on plant interactions mediated by the biophysical dynamics of a shared soil resource, accounting for competition for a shared limiting resource and the positive effects of ecosystem engineers on their neighbors due to resource mining. Resource mining is just one possible mechanism among many leading to positive effects that one plant can have over its neighbors, and it may lead to facilitation if such positive effects overcome competition. Based on this particular mechanism, our results show that, at least in theory, antagonistic facilitation between pairs of plants can be evolutionarily stable in stressful environments even when the nurse plant incurs in energetic costs. However, in practice this result is limited by the particular factor allowing facilitation, regardless of whether nurse plants incur in a cost to facilitate their neighbors or not. It is still uncertain whether antagonistic facilitation may or may not be fixed by evolution when nurse plants protect their neighbors from herbivory, cast shade over them or act as physical barriers baffling strong winds or buffering extreme temperatures, to name a few alternative mechanisms that have been suggested as forces of facilitation in experimental studies. Moreover, even when resource engineering drives facilitation, relaxing some of the model assumptions considered here could also change the conditions in which antagonistic facilitation is predicted in theory. For example, plant species might differ in their resource uptake strategies, defining tradeoffs in resource uptake that might alter the stability conditions for antagonistic facilitation. Alternatively, environmental gradients may be defined by several parameters changing simultaneously, even in opposite directions. Lastly, larger population sizes with different compositions and spatial structures could also change the conditions in which facilitation via soil resource engineering is evolutionarily stable. Here, we provide a first step to better understand how environmental conditions and their underpinning biophysical mechanisms might make antagonistic facilitation evolutionarily stable. However, more work is still needed to investigate the evolutionary stability of antagonistic facilitation based on other mechanisms and across environmental gradients.

## Acknowledgments

This work was partially funded by the Center of Advanced Systems Understanding (CASUS) which is financed by Germany’s Federal Ministry of Education and Research (BMBF) and by the Saxon Ministry for Science, Culture and Tourism (SMWK) with tax funds on the basis of the budget approved by the Saxon State Parliament. GAM was supported by a Postdoctoral Fellowship from São Paulo Research Foundation (FAPESP), grant 2019/21227-0. RMG was partially supported by the Simons Foundation through grant 284558FY19; FAPESP through ICTP-SAIFR grant 2021/14335-0 and the Jovem Pesquisador grant 2019/05523-8. This research was supported by resources supplied by the Center for Scientific Computing (NCC/GridUNESP) of the São Paulo State University (UNESP).

## Author contributions

Ciro Cabal and Ricardo Martinez-Garcia conceived the project; Gabriel A. Maciel, Ciro Cabal and Ricardo Martinez-Garcia designed the model; Gabriel A. Maciel led model analysis with input from Ricardo Martinez-Garcia and Ciro Cabal; Ciro Cabal led manuscript writing with significant contributions from Gabriel A. Maciel and Ricardo Martinez-Garcia; Ciro Cabal led visualization with contributions from Gabriel A. Maciel and Ricardo Martinez-Garcia. All authors gave final approval for paper publication.

## Data accessibility statement

All the Python scripts with model implementation are available at https://github.com/gandreguetto/antagonistic-facilitation.

