## Supplementary material for "The evolutionary stability of plant antagonistic facilitation across environmental gradients and its ecological consequences: soil resource engineering as a case study": Fig. SM1

### Figure SM1:

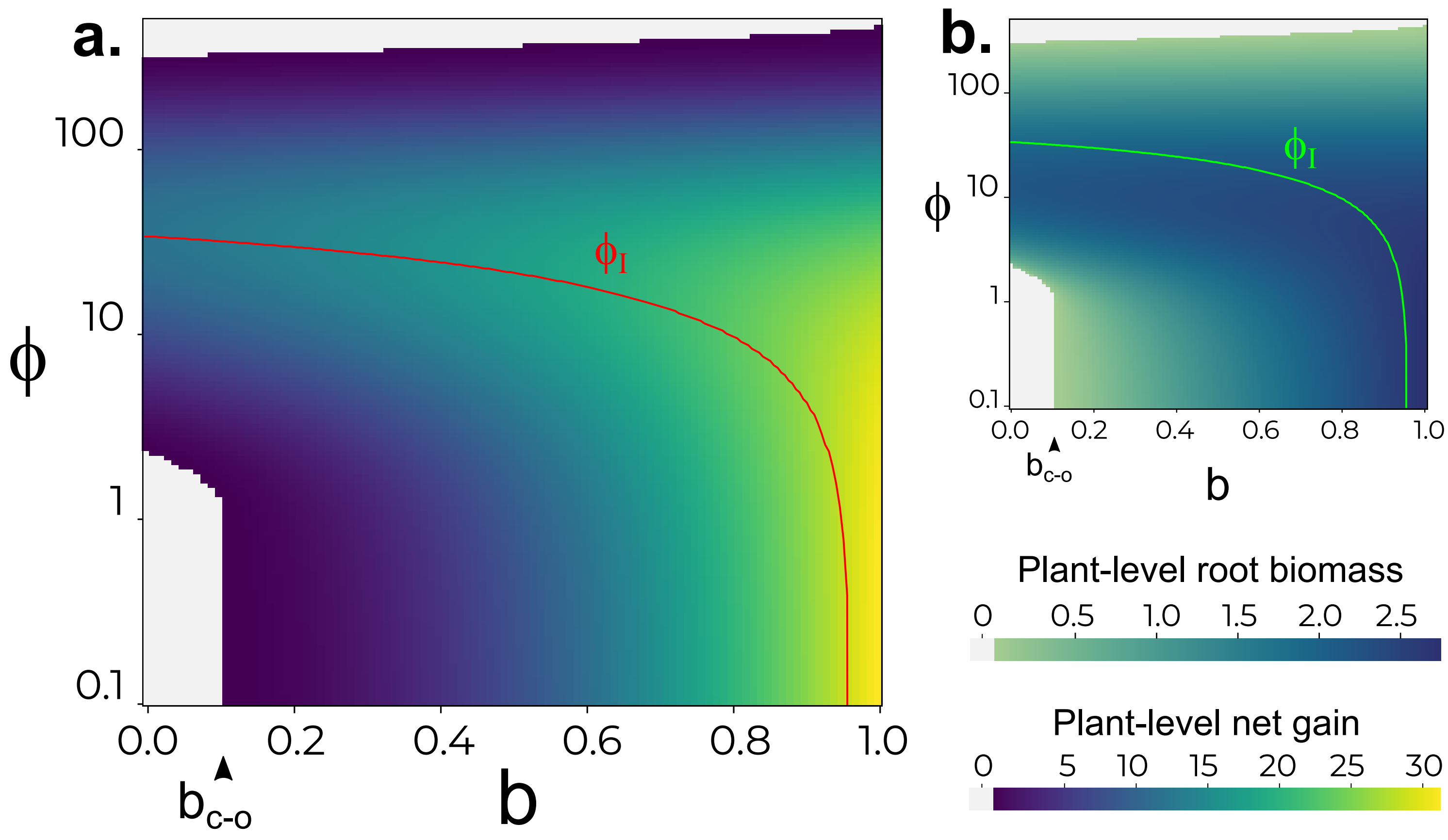


**CAPTION: a-** Plant-level net gain and **b-** fine root biomass as a function of $\phi$ (log-scaled axis) and of *b* for an ecosystem engineer growing alone. Lines in both panels represent the value of the mining trait that evolves at different successional times, $\phi_I$. The point $b_{c\text{-}o}$ is the stress threshold below which opportunistic plants cannot survive (light gray region).
