## Supplementary material for "The evolutionary stability of plant antagonistic facilitation across environmental gradients and its ecological consequences: soil resource engineering as a case study": Fig. SM2

### Figure SM2:

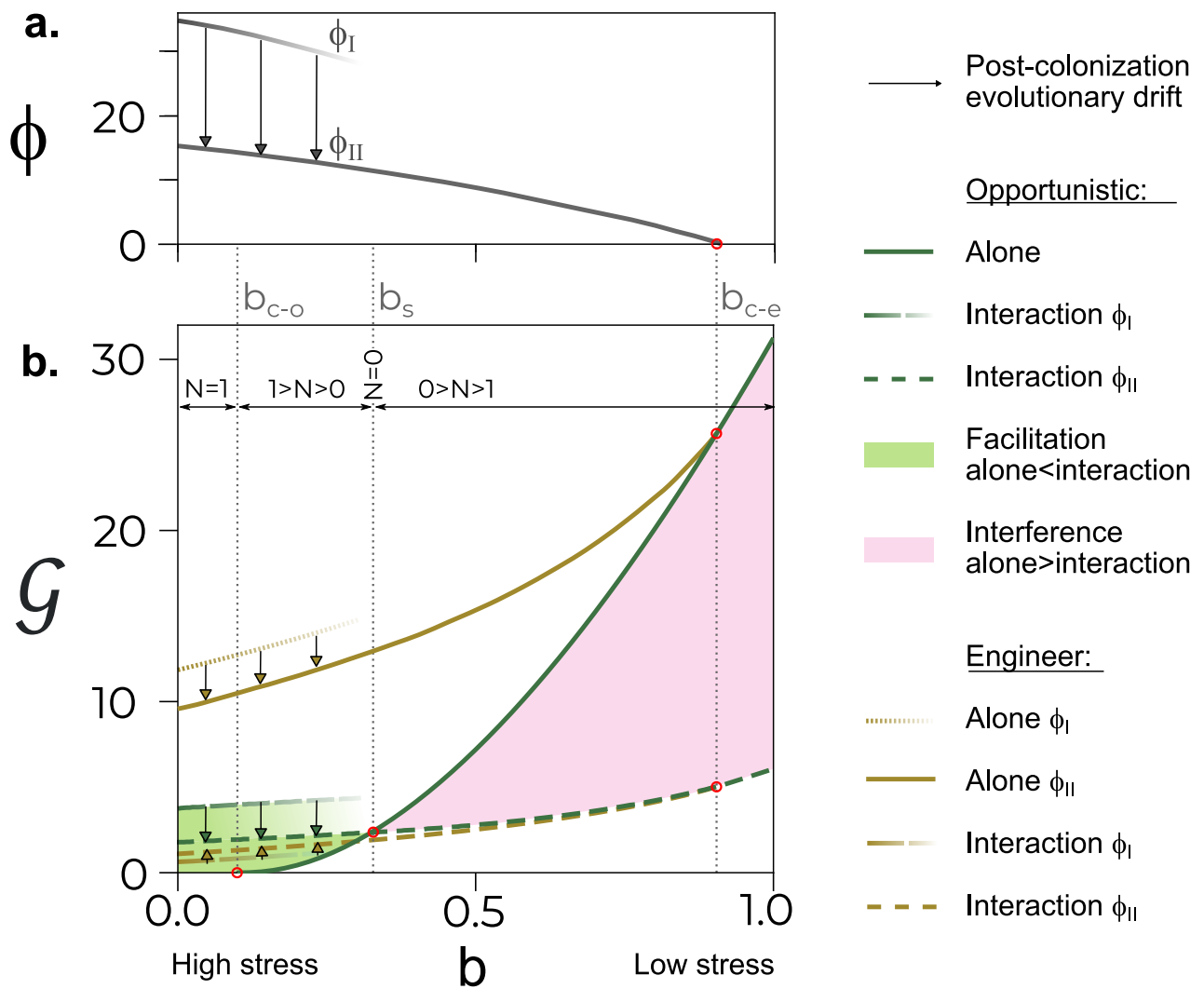

**CAPTION: a-** Mining trait $\phi_I$ in an evolutionary equilibrium resulting from the model parameterization step of the analysis across an environmental gradient defined by $b$ (proportion of the resource spontaneously available). **b-** Plant-level net resource gain ($\mathcal{G}$) as a function of $b$ for plants growing alone and engineer-opportunistic pairs. Different lines show different evolutionary equilibria as indicated in the legend: $\phi_I$ (only shown for low-$b$ scenarios where engineer plants might evolve alone) and $\phi_{II}$.

Figure 2 in the main text is a schematic summary of the results from this figure, in which $RII_G$ represents the normalized difference between values of $G$ shown in this figure comparing opportunistic plants alone vs. interacting with an ecosystem engineer.
