## Supplementary material for "The evolutionary stability of plant antagonistic facilitation across environmental gradients and its ecological consequences: soil resource engineering as a case study": Fig. SM3

### Figure SM3:

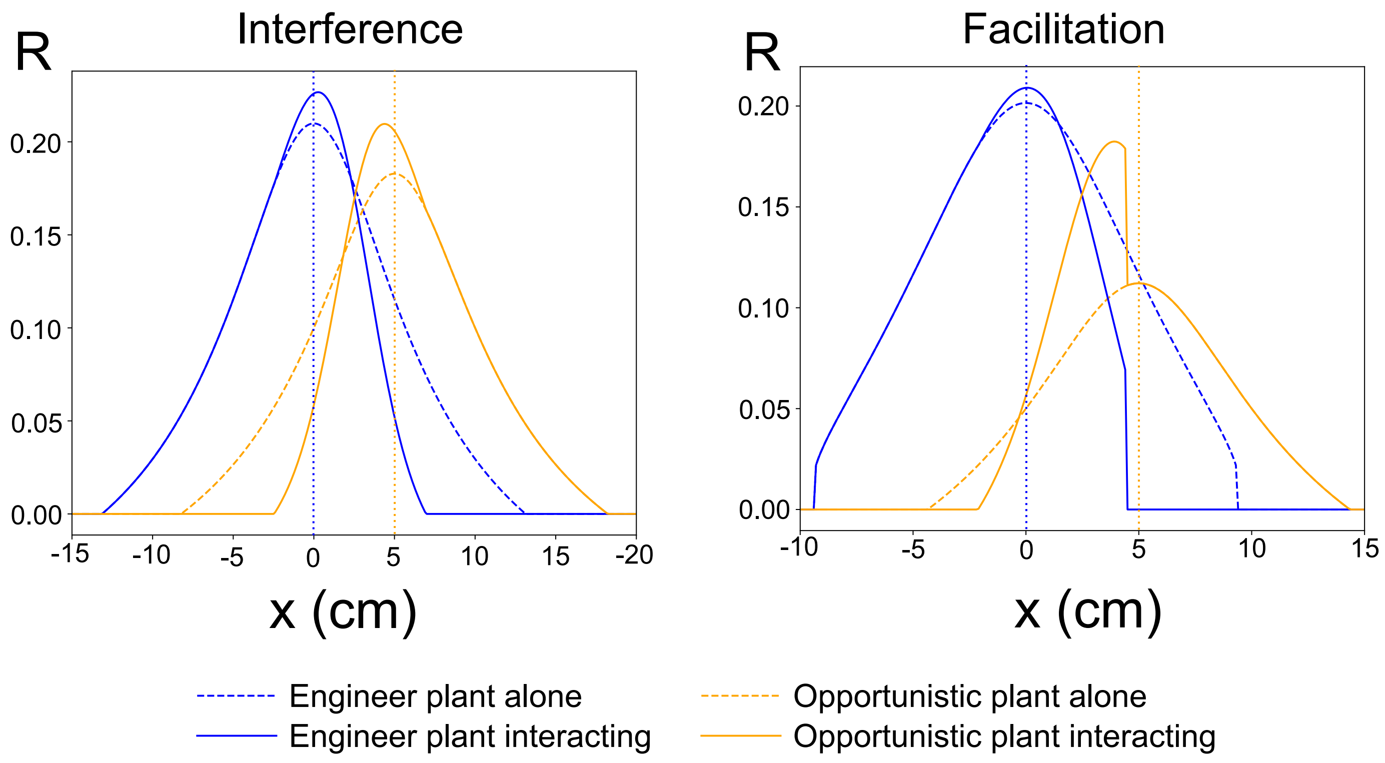


**CAPTION:** Root density distribution of ecosystem engineer (blue) centered in x = 0 cm and opportunistic plants (orange) centered in x = 5 cm, growing alone (dashed lines) or interacting with d = 5 cm (solid lines). Left figure represents a case of interference (for b = 0.8 we have $RII_G$ = - 0.3060) and right figure a case of facilitation (for b = 0.45 we have $RII_G$ = 0.0107). When comparing how the foraging behavior of the opportunistic plant changes from growing alone to interacting we see similar results whether *b* leads to facilitation or interference; root density increases near the insertion point at x = 5 cm, skewing the root allocation towards the neighbor, but the root range diminishes. Therefore, observing this root density distribution patterns cannot be claimed to prove whether the interaction outcome is facilitation or interference. Other parameter values are Cb = 5.0, Ct = 0.2, Ce = 0.1, $\delta^{-1}$ = 0.1.
