## Supplementary material for "The evolutionary stability of plant antagonistic facilitation across environmental gradients and its ecological consequences: soil resource engineering as a case study": SM Text

**2 Model experiments** **7**

*These authors contributed equally to this work and share first authorship
†

In this document, we detail the two-step model analysis. In section 1 we describe the calculation of the values of the resource-mining trait used in the simulated experiments, which are explained in detail in Section 2. Throughout the text, we also provide various figures showing the spatial distribution of root biomass predicted by the model in the different scenarios we cover.

### 1 Model parameterization

#### 1.1 Evolution of resource mining in solitary soil-engineer plants.

Departing from Eqs. (2)-(4) in the main text, we can write the resource dynamics and net resource gain of a solitary soil engineer as:

$$\frac{\partial W(\ell,\,t)}{\partial t} = I(\phi, R_{\rm E}) - \delta\, W(\ell,t) - \alpha\, W(\ell,t) R_{\rm E}(\ell,t) \tag{S1.1}$$

$$G_{\rm E}(\ell,t) = \Big[\mathrm{WUE}\,\alpha\, W(\ell,t) - C(\ell,\,\phi)\Big] R_{\rm E}(\ell,t). \tag{S1.2}$$

where $I(\phi, R_{\rm E})$ is the monotonically increasing and saturating function of $\phi R_{\rm E}$ introduced in Eq. (1) of the main text and $C(\ell,\phi)$ is the cost function in Eq. (4) of the main text.

If we further assume that resource dynamics is much faster than plant growth, we can consider that the resource is always at equilibrium and Eq. (S1.1) reduces to

$$W^*(\ell) = \frac{I(\phi, R_{\rm E})}{\delta + \alpha R_{\rm E}(\ell)} \tag{S1.3}$$

Substituting Eq. (S1.3) into (S1.2) we arrive to Eq. (5) of the main text particularized for a solitary soil-engineer plant

$$G_{\rm E}(\ell,t) = \left[\frac{\mathrm{WUE}\,\alpha\, I(\phi,R_{\rm E})}{\delta + \alpha R_{\rm E}(\ell)} - C(\ell,\phi)\right] R^*_{\rm E}(\ell,t). \tag{S1.4}$$

Next, we obtain the spatial distribution of root biomass and the value of the resource-mining trait, $R^*_{\rm E}(\ell)$ and $\phi_{\rm I}$ respectively, that maximize the net resource-gain function in Eq. (S1.4). First, we explain the steps to obtain $R^*_{\rm E}(\ell)$ assuming that $\phi$ is a free parameter and can take any value. Then, we explain the case in which the plant evolves a resource-mining trait value $\phi = \phi_I$ and changes its spatial distribution of roots according to the new mining intensity.

##### 1.1.1 Calculation of the optimal spatial distribution of roots assuming $\phi$ as a free parameter

To obtain $R^*_{\rm E}$ we impose a maximization condition in Eq. (S1.4)

$$\frac{\partial G}{\partial R_{\rm E}} = 0 \tag{S1.5}$$

that, after making explicit the functional form of $I(\phi, R_{\rm E})$, leads to the following fourth-order polynomial equation for $R_{\rm E}$,

$$\sum_{i=0}^{4} A_i R^i_{\rm E} = 0 \tag{S1.6}$$

where

$$A_4 = \phi^2 \tag{S1.7}$$

$$A_3 = 2\phi\left(1 + \frac{\delta}{\alpha}\phi\right) \tag{S1.8}$$

$$A_2 = \left(1 + \frac{\omega \mathrm{WUE}}{C(\ell,\phi)}\phi\, b\right) + \left(4\frac{\delta}{\alpha} - \frac{\omega\mathrm{WUE}}{C(\ell,\phi)}\right)\phi + \left(\frac{\delta}{\alpha} - \frac{\omega\mathrm{WUE}}{C(\ell,\phi)}\right)\frac{\delta}{\alpha}\phi^2 \tag{S1.9}$$

$$A_1 = 2\frac{\delta}{\alpha}\left[1+\left(\frac{\delta}{\alpha}-\frac{\omega \mathrm{WUE}}{C(\ell,\phi)}\right)\phi\right] \tag{S1.10}$$

$$A_0 = \frac{\delta}{\alpha}\left(\frac{\delta}{\alpha}-\frac{\omega \mathrm{WUE}}{C(\ell,\phi)}b\right) \tag{S1.11}$$

We calculate the value of the engineer root biomass density that results in the global maximum of the net resource-gain function in two steps. First, we solve Eq. (S1.6) numerically. Second, from all the real positive roots such that $G(R_{\mathrm{E}}) > 0$, $R^*_{\mathrm{E}}$ is the one that gives the largest $G$. If Eq. (S1.6) has no positive real roots or none of them results in a positive net resource gain, the plant does not grow roots at that soil location, and $R^*_{\mathrm{E}}(\ell) = 0$.

From the coefficients in Eqs. (S1.7)-(S1.11), Eq. (S1.6) has real and positive solutions only if

$$\frac{\omega\,\alpha\,\mathrm{WUE}}{\delta} > C(\ell,\phi). \tag{S1.12}$$

That is, a necessary condition for plant growth is that the product of the uptake rate per unit of fine root mass, the potential resource input, and the resource use efficiency must be larger than the root maintenance and production cost times the resource abiotic decay rate. Otherwise, all the coefficients in Eqs. (S1.7)-(S1.11) are positive, Eq. (S1.6) never changes in sign and the net resource-gain has no strictly positive extrema.

Because the cost function $C(\ell,\phi)$ increases with increasing $\ell$ and all other parameters in the inequality (S1.12) are constant, (S1.12) indicates that the plant root system must be finite. Moreover, the extension of the root system $\ell_c$ is defined by the first value of $\ell$ at which the global maximum of the net resource-gain function shifts from $R^*_{\mathrm{E}} > 0$ to $R^*_{\mathrm{E}} = 0$ (see Cabal et al. 2020 for a detailed calculation of these properties of the root system in a simpler scenario).

Finally, once we obtain the engineer root density that maximizes the net resource gain at each soil location, we can obtain the plant-level root biomass by integrating the root biomass density over the extension of the plant root system,

$$\mathcal{R} = \int_{-\ell_c}^{\ell_c} R^*_{\mathrm{E}}(\ell)d\ell \tag{S1.13}$$

Similarly, we can calculate the plant-level net resource gain by integrating the net resource-gain function over the entire root system,

$$\mathcal{G} = \int_{-\ell_c}^{\ell_c} G_{\mathrm{E}}(R^*_{\mathrm{E}}(\ell),\ell)d\ell \tag{S1.14}$$

These two quantities in Eqs. (S1.13)-(S1.14) are shown in Fig. 2 of the main text. Here we show in Fig. S1 the spatial distribution of root density predicted by our model for different values of the fraction of resource spontaneously available $b$ and the resource mining trait $\phi$. In panel (A) we choose low $\phi$ and $b > b_{\text{c-o}}$. In this limit, resource mining alone can not sustain the plant, but it can grow without mining resources because there is a large fraction of resources spontaneously available. Panel B shows the opposite extreme $b < b_{\text{c-o}}$ and intermediate $\phi$. In this limit, the plant needs to mine resources to grow, and $\phi$ is within the range of intermediate values that allow plant growth. In this regime, we observe a qualitative change in the shape of the spatial distribution of roots. The root biomass density drops sharply at a given distance to the plant insertion point. This abrupt decline in root density appears because plant survival depends on mining resources, which in turn depends on root density. Past some distance to the insertion point, the root biomass density is so low that roots cannot mine enough resources and the root system ends abruptly. Finally, panel C shows a root profile for $b > b_{\text{c-o}}$ and intermediate $\phi$. In this regime, resource mining is not necessary for survival, but it still contributes to enhanced plant growth [see the main text for a longer discussion of these three regimes in the $(b,\,\phi)$ parameter space].

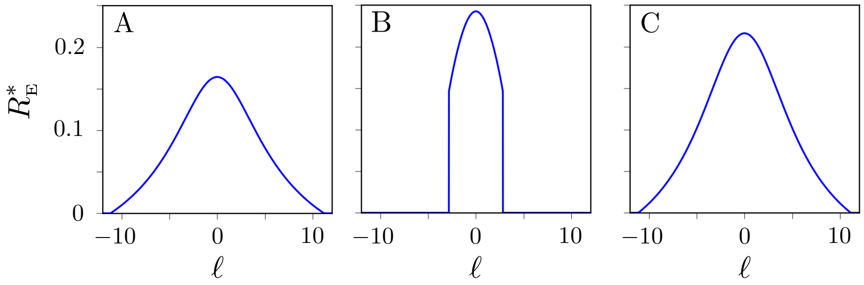

Fig. S1: Root distributions for three different values of the resource mining trait. (A) $\phi = 0.5$ and $b = 0.6$. (B) $b = 0.05$ and $\phi = 3$. (C) $b = 0.6$ and $\phi = 0.3$. Other parameters are: $\omega = 5$, WUE $= 1$ $\delta = 0.1$, $\alpha = 1$, $C_b = 5$, $C_t = 0.2$ and $C_e = 0.1$.

##### 1.1.2 Calculation of optimal mining intensity and spatial distribution of roots, $(\phi_{\text{I}}, R^*_{\mathbf{E}})$.

To perform this calculation, we repeat the procedure described in Section 1.1.1 for several values of $\phi$, leading to a spatial distribution of root biomass for each value of $\phi$. Finally, we obtain the evolved value of the resource-mining trait $\phi_{\text{I}}$ as the value of $\phi$ that results in larger plant-level net resource gain, $\mathcal{G}$. The predicted spatial distribution of root biomass will be the one associated with that specific $\phi$ (see Fig. S3 for examples of how root systems change when $\phi$ evolves to $\phi_{\text{I}}$ as opposed to opportunistic plants with $\phi = 0$). If $\phi_{\text{I}} \neq 0$ for a given parameterization, the model predicts soil engineering to evolve in the environmental conditions represented by that set of parameters. Otherwise, the model predicts resource mining to be evolutionarily unstable. This value of $\phi_{\text{I}}$ is shown in the main text in both panels of Fig. 2 (red and green lines overlaid on the density plots).

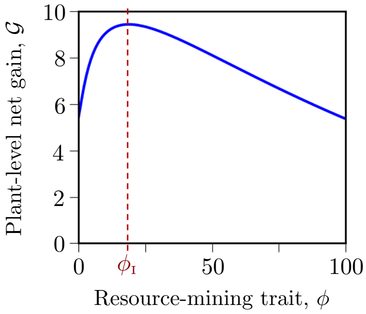

Fig. S2: Plant-level net gain as defined in Eq. (S1.14) as a function of the resource-mining trait $\phi$. To obtain $\mathcal{G}$ for each $\phi$ we first obtain the spatial distribution of root biomass that maximizes the net resource-gain function. Model parameters: $b = 0.6$, WUE $= 1$, $\omega = 5$, $\alpha = 1$, $\delta = 0.1$, $C_b = 5$, $C_t = 0.2$ and $C_e = 0.1$.

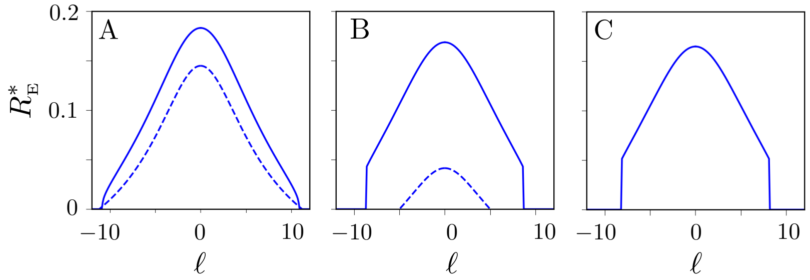

Fig. S3: Spatial distribution of roots for different values of $b$. (A) $b = 0.6$, (B) $b = 0.2$ and (C) $b = 0.05$. Dashed curves show model predictions in the absence of resource mining ($\phi = 0$). Solid lines show the root distribution when soil-engineer plants evolve the resource mining trait isolated from any other individual: (A) $\phi_{\text{I}} = 19$ (corresponding to the maximum of the curve in Fig. S2), (B) $\phi_{\text{I}} = 30.5$, and (C) $\phi_{\text{I}} = 34$. Other parameter values: WUE $= 1$, $\omega = 5$, $\alpha = 1$, $\delta = 0.1$, $C_b = 5$, $C_t = 0.2$ and $C_e = 0.1$.

#### 1.2 Evolution of resource mining in a soil-engineer plant interacting with a spreading opportunistic. Calculation of $\phi_{\text{II}}$

In the main text, we investigate the nature of the interaction between a soil engineer and an opportunistic plant (Sections 3.2 and 3.3 of the main text). To conduct this analysis, we first mimic the evolutionary dynamics of the engineer plant in a situation in which it grows within a background of spreading opportunistic plants. In this section of the Supplementary Text, we provide details on how to obtain the evolutionarily stable solutions of this interaction between a soil engineer and a spreading opportunistic plant.

First, we need to extend the model equations to include the effect of spreading opportunistic plants. If we assume that both types of plants are identical except in their ability to mine resources, all parameters are plant-type independent. The resource dynamics is

$$\frac{\partial W(\ell,t)}{\partial t} = I(\phi, R_{\text{E}}) - \delta W(\ell,t) - \alpha\Big[R_{\text{E}}(\ell,t) + R_{\text{O}}(\ell,t)\Big]W(\ell,t) \tag{S1.15}$$

where the subscripts E and O indicate engineer and opportunistic, respectively. The net resource-gain function for each plant type is

$$G_{\text{E}}(\ell,t) = \Big[\text{WUE}\,\alpha\,W(\ell,t) - \Big(c_b + c_t\ell^2 + c_e\phi\Big)\Big]R_{\text{E}}(\ell,t) \tag{S1.16}$$

$$G_{\text{O}}(\ell,t) = \Big[\text{WUE}\,\alpha\,W(\ell,t) - c_b\Big]R_{\text{O}}(\ell,t) \tag{S1.17}$$

where we have already considered that opportunistic plants are spreading plants and thus $c_t = 0$ (see main text), and they do not incur any resource-mining cost $c_e = 0$ or $\phi = 0$. Following the same time scale separation we introduced for solitary plants, we can assume that the resource concentration is always at equilibrium,

$$W^*(\ell) = \frac{I(R_{\text{E}},\phi)}{\delta + \alpha[R_{\text{E}}(\ell) + R_{\text{O}}(\ell)]} \tag{S1.18}$$

and insert this expression in Eqs. (S1.16)-(S1.17) to eliminate the dependence on $W$ and obtain a closed system of equations for $R_{\text{E}}$ and $R_{\text{O}}$.

First, we consider that the spatial distributions of roots changes because of the engineer-opportunistic interaction, but the engineer resource-mining trait remains constant and equal to the value evolved in the absence of spreading opportunistic individuals, $\phi_{\text{I}}$. To obtain the new spatial configuration of root densities, we first calculate the density of roots for each plant in a hypothetical coexistence $R_{\text{E}}(\ell) > 0$ and $R_{\text{O}}(\ell) > 0$ state solving

$$\frac{\partial G_{\text{E}}}{\partial R_{\text{E}}} = 0, \tag{S1.19}$$

$$\frac{\partial G_{\text{O}}}{\partial R_{\text{O}}} = 0. \tag{S1.20}$$

To this end, we first use the equation that results from solving Eq. (S1.20) to obtain $R_{\text{O}}$ as a function of $R_{\text{E}}$:

$$R_{\text{O}} = \sqrt{\frac{\text{WUE}}{c_b}\,I(\phi,R_{\text{E}})\left(\frac{\delta}{\alpha} + R_{\text{E}}\right)} - \left(\frac{\delta}{\alpha} + R_{\text{E}}\right). \tag{S1.21}$$

Next, we insert Eq. (S1.21) in Eq.(S1.19) to get a $7th$-order polynomial equation in $R_{\text{E}}$ that we solve numerically to obtain the positive real solutions for $R_{\text{E}}(\ell)$ and $R_{\text{O}}(\ell)$.

Once we calculate the density of roots in a hypothetical coexistence state solving Eqs. (S1.19)-(S1.20), we replace the values of $R_{\text{E}}(\ell)$ and $R_{\text{O}}(\ell)$ in Eqs. (S1.16)-(S1.17) and evalute the net resource intake of each plant. If both plants have a positive net resource intake in the coexistence state, we choose this solution as $R^*_{\text{E}}(\ell)$ and $R^*_{\text{O}}(\ell)$. If both plants have negative net-resource intake, none of the plants can grow resources in that soil coordinate and $R^*_{\text{E}}(\ell) = R^*_{\text{O}}(\ell) = 0$. We find this scenario in highly stressed environments ($b$

low), where resource mining is obligatory, and in soil coordinates far from the engineer insertion point, where roots are expensive to produce. Finally, if $G_{\mathrm{E}} < 0$ and $G_{\mathrm{O}} > 0$ only the opportunistic plant grows roots and $R^*_{\mathrm{E}}(\ell) = 0$, $R^*_{\mathrm{O}}(\ell) \neq 0$. We calculate the root biomass of the opportunistic plant following the steps in Section 1.1.1 with $\phi = 0$. We find this scenario in mildly stressed environments ($b$ high), where resource mining is facultative, and in soil coordinates far from the engineer insertion point, where roots are expensive to produce. See Fig. S4 for examples of spatial distributions of root and net resource intake.

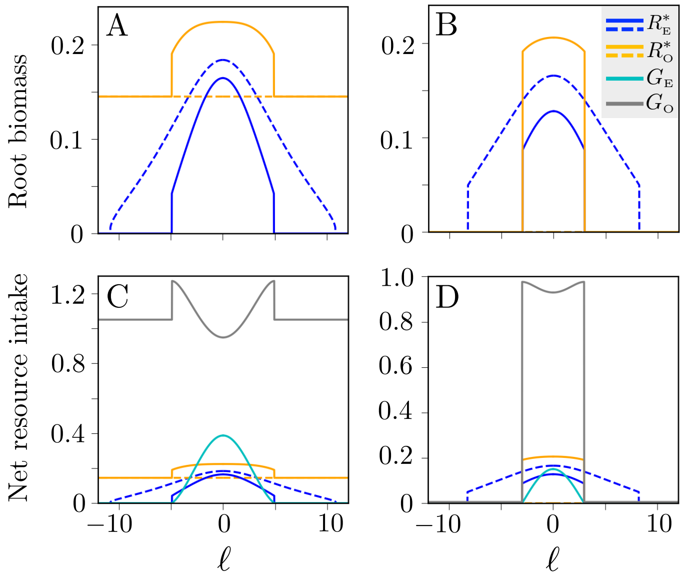

Fig. S4: Root distribution (A, B) and net resource intake (C, D) for two different levels of environmental stress and assuming that the resource mining trait evolved in solo plants, $\phi = \phi_{\mathrm{I}}$. (A, C) Represent a low stress level in which the opportunistic plant can survive alone, $b = 0.6$ and $\phi_{\mathrm{I}} = 18.8$. (B, D) Represent a high stress level in which resource mining is obligatory, $b = 0.05$ and $\phi_{\mathrm{I}} = 34$. In the root density panels, the solid lines correspond to a soil engineer (blue) and a spreading opportunistic (orange) plant in interaction. The dashed lines show the expected distribution of roots for the same individuals when they grow alone. Other parameter values: WUE $= 1$, $\omega = 5$, $\delta = 0.1$, $\alpha = 1$, $c_b = 5$, $c_t = 0.2$, and $c_e = 0.1$.

Finally, we allow the resource-mining trait to evolve following the interaction between the soil engineer and the opportunistic spreading plant. To this end, we extend the protocol introduced in Section 1.1.2 to the case where there are two plants and, thus, two net resource-gain functions. First, assuming an arbitrary $\phi$, we obtain the engineer and spreading opportunistic root systems that jointly maximize each plant-level net resource gain. Second, we repeat this calculation for several values of $\phi$. Finally, we obtain $\phi_{\mathrm{II}}$ as the value of $\phi$ for which the associated root systems result in a maximum plant-level net gain. This value of $\phi_{\mathrm{II}}$ is shown in the top panel of Fig. 3 in the main text [see Fig. S5 for a comparison between the root systems of engineer and opportunistic plants in interaction ($\phi = \phi_{\mathrm{II}}$ and $\phi = \phi_{\mathrm{I}}$) and growing solo].

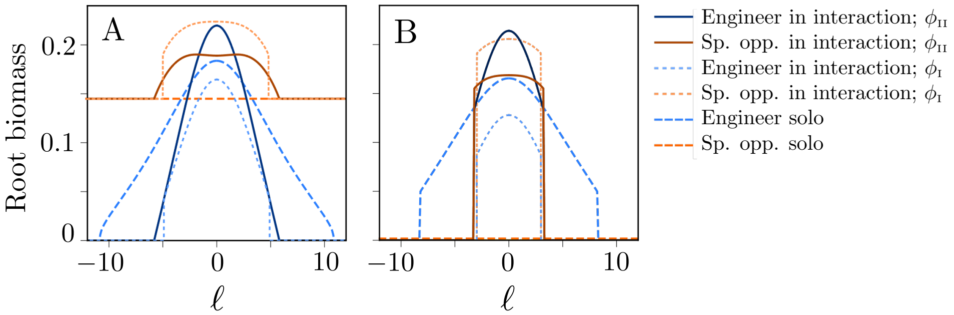

Fig. S5: Root distribution for $b = 0.6$ (A) and $b = 0.05$ (B). Solid and dashed lines of different colors follow the figure legend. Curves with resource-mining evolving in solitary engineer plants have the same values that in Fig. S4: $\phi_{\mathrm{I}} = 18.8$ (A) and $\phi_{\mathrm{I}} = 34$ (B). Curves using the resource-mining trait value evolved in interacting plants have $\phi_{\mathrm{II}} = 5$ (A) and $\phi_{\mathrm{II}} = 12$ (B). Other parameters: WUE $= 1$, $\omega = 5$, $\delta = 0.1$, $\alpha = 1$, $c_b = 5$, $c_t = 0.2$, and $c_e = 0.1$.

#### 2 Model experiments

To perform the simulated experiments for the interaction between the soil engineer and normal opportunistic plants, we first need to modify the model equations to account for the fact that the normal opportunistic builds thick roots

$$G_{\mathrm{E}}(\ell,t) = \left[\mathrm{WUE}\,\alpha\,W(\ell,t) - \left(c_b + c_t\ell^2 + c_e\phi\right)\right] R_{\mathrm{E}}(\ell,t) \tag{S2.1}$$

$$G_{\mathrm{O}}(\ell,t) = \left[\mathrm{WUE}\,\alpha\,W(\ell,t) - c_b - c_t(\ell-d)^2\right] R_{\mathrm{O}}(\ell,t) \tag{S2.2}$$

The resource dynamics is given by Eq. (S1.15). For the calculations in the main text, we choose $d = 0$ in Eq. (S2.2) to maximize the interaction strength, but also consider $d = 5$ in Fig. SM3. For the resource-mining trait (in Figure 3 of the main text), we use $\phi = \phi_{\mathrm{I}}$ or $\phi_{\mathrm{II}}$ depending on whether we are interested in considering a soil-engineer plant that evolved resource mining isolated or in interaction with an opportunistic spreading plant. In Figure 4, we assumed that resource-mining evolved in the presence of opportunistic plants and therefore used $\phi = \phi_{\mathrm{II}}$) for each of the environmental conditions we considered.

To calculate the predicted spatial distribution of roots for the soil engineer and the opportunistic plant, we follow the steps summarized in Eqs. (S1.18)-(S1.21) to obtain a seventh-order polynomial equation in $R_{\mathrm{E}}$. We solve this equation numerically to get $R^*_{\mathrm{E}}$ as the real positive solution maximizing $\mathcal{G}_{\mathrm{E}}$ and replace these solutions in Eq. (S1.21) to obtain $R^*_{\mathrm{O}}$. If this solution exists, is real positive, and leads to positive plant-level resource gain, we consider that the spreading opportunistic and the engineer can coexist. Otherwise, if the solution does not exist or leads to non-positive plant-level net gain, the interaction results in the competitive exclusion of the normal opportunistic plant.
